# SCRAPP: A tool to assess the diversity of microbial samples from phylogenetic placements

**DOI:** 10.1101/2020.02.28.969980

**Authors:** Pierre Barbera, Lucas Czech, Sarah Lutterop, Alexandros Stamatakis

**Author notes:** To whom correspondence should be addressed. E-mail: {, }.

## Abstract

Microbial ecology research is currently driven by the continuously decreasing cost of DNA sequencing and the improving accuracy of data analysis methods. One such analysis method is phylogenetic placement, which establishes the phylogenetic identity of the anonymous environmental sequences in a sample by means of a given phylogenetic reference tree. However, assessing the diversity of a sample remains challenging, as traditional methods do not scale well with the increasing data volumes and/or do not leverage the phylogenetic placement information.

Here, we present `SCRAPP`, a highly parallel and scalable tool that uses a molecular species delimitation algorithm to quantify the diversity distribution over the reference phylogeny for a given phylogenetic placement of the sample. `SCRAPP` employs a novel approach to cluster phylogenetic placements, called placement space clustering, to efficiently perform dimensionality reduction, so as to scale on large data volumes. Furthermore, it utilizes the phylogeny-aware molecular species delimitation method `mPTP` to quantify diversity.

We evaluated `SCRAPP` using both, simulated and empirical datasets. We use simulated data to verify our approach. Tests on an empirical dataset show that SCRAPP-derived metrics can classify samples by their diversity-correlated features equally well or better than existing, commonly used approaches.

`SCRAPP` is available at https://github.com/pbdas/scrapp

## 1 INTRODUCTION

Environmental microbial DNA sampling is increasingly becoming a standard practice, not least due to continuously decreasing sequencing costs. One, by now, established way to analyze such data is Phylogenetic Placement (Berger et al., 2011; Matsen et al., 2010; Barbera et al., 2018). In phylogenetic placement (hereafter called placement), sequences from environmental samples (query sequences, QS) are placed on a phylogenetic tree comprising the biome under study (henceforth denoted as reference tree), resulting in a set of QS placements on this reference tree (hereafter called placements). Placement has, for instance, been successfully applied to describe the composition of a soil protist environment (Mahé et al., 2017) or to identify the relationships between bacterial community composition and disease (Srinivasan et al., 2012).

Another key goal of molecular microbial studies is to assess microbial diversity. A plethora of distinct metrics already exist to quantify the diversity of a sample, or parts thereof (Tucker et al., 2017). Of these metrics, perhaps the most widely used ones rely on phylogenetic information (e.g., the UniFrac distance (Lozupone & Knight, 2005) or the Phylogenetic Diversity (PD) measure (Faith, 1992)). A relatively recent approach to quantifying diversity using sequence data is phylogeny-aware molecular species delimitation (Fujisawa & Barraclough, 2013; Yang, 2015; Zhang et al., 2013; Kapli et al., 2017). These methods rely on a given phylogenetic tree to identify species boundaries, essentially resulting in a clustering of the tips into distinct species.

Here, we combine previous work on phylogenetic placement (Barbera et al., 2018) and species delimitation (Zhang et al., 2013; Kapli et al., 2017) to devise a measure of phylogeny-aware relative within-sample diversity. Our `SCRAPP` (Species Counting on Reference trees viA Phylogenetic Placement) tool, quantifies diversity by initially grouping placements (QS) by the branch on the reference tree (reference branch) to which they most likely belong with respect to their phylogenetic likelihood score. Subsequently, for each such group of QS placed onto the same reference branch, we infer a separate phylogenetic tree comprising the QS of that group, optionally including an outgroup sequence from the reference tree. We call such a tree a branch query phylogeny (BQP). Finally, we apply `mPTP` (Kapli et al., 2017) to the BQP to obtain a species count for the corresponding reference branch. The output of `SCRAPP` is a branch-annotated reference tree that depicts how species diversity is distributed over the reference tree. `SCRAPP` is implemented in Python and relies on `mpi4py` (Dalcín et al., 2005, 2008; Dalcin et al., 2011) for the respective parallel implementation targeting both, shared, and distributed memory systems.

Some concepts are based on our admittedly difficult to use EPA-PTP tool, an early attempt to integrate placement with species delimitation (Zhang et al., 2013). The goal of `SCRAPP` is thus to quantify diversity for each branch of the reference tree individually and to improve usability. In contrast to `SCRAPP`, EPA-PTP used placement to calculate a single, overall species delimitation over the entire reference tree extended by all BQPs simultaneously.

## 2 DESCRIPTION

In the following, we initially provide a detailed description of the `SCRAPP` tool. An overview is provided in Figure 1. `SCRAPP` takes as input a `jplace` (Matsen et al., 2012) file containing the placements and the associated reference tree, as well as the corresponding multiple sequence alignment (MSA) of the QS. From this, we generate per-branch QS MSAs. These include all QS whose most likely placement was on the given branch. However, we remove those placements from this set, whose best likelihood weight ratio (LWR) (von Mering et al., 2007) is below a given threshold (`--min-weight`, default 0.5).

If desired, an outgroup from a user specified reference MSA is included in each branch QS MSA such that the corresponding BQP that is produced in the subsequent step can be rooted at this outgroup. We automatically choose the outgroup sequence for a given BQP as the leaf sequence in the reference tree that is most distant from the given branch. Note that, `mPTP` species delimitation operates on rooted phylogenies. Thus, specifying an outgroup can be beneficial if a more reliable root for the BQP is desired. If a root is not provided, `mPTP` will automatically root the BQP on its longest branch.

**Figure 1:**
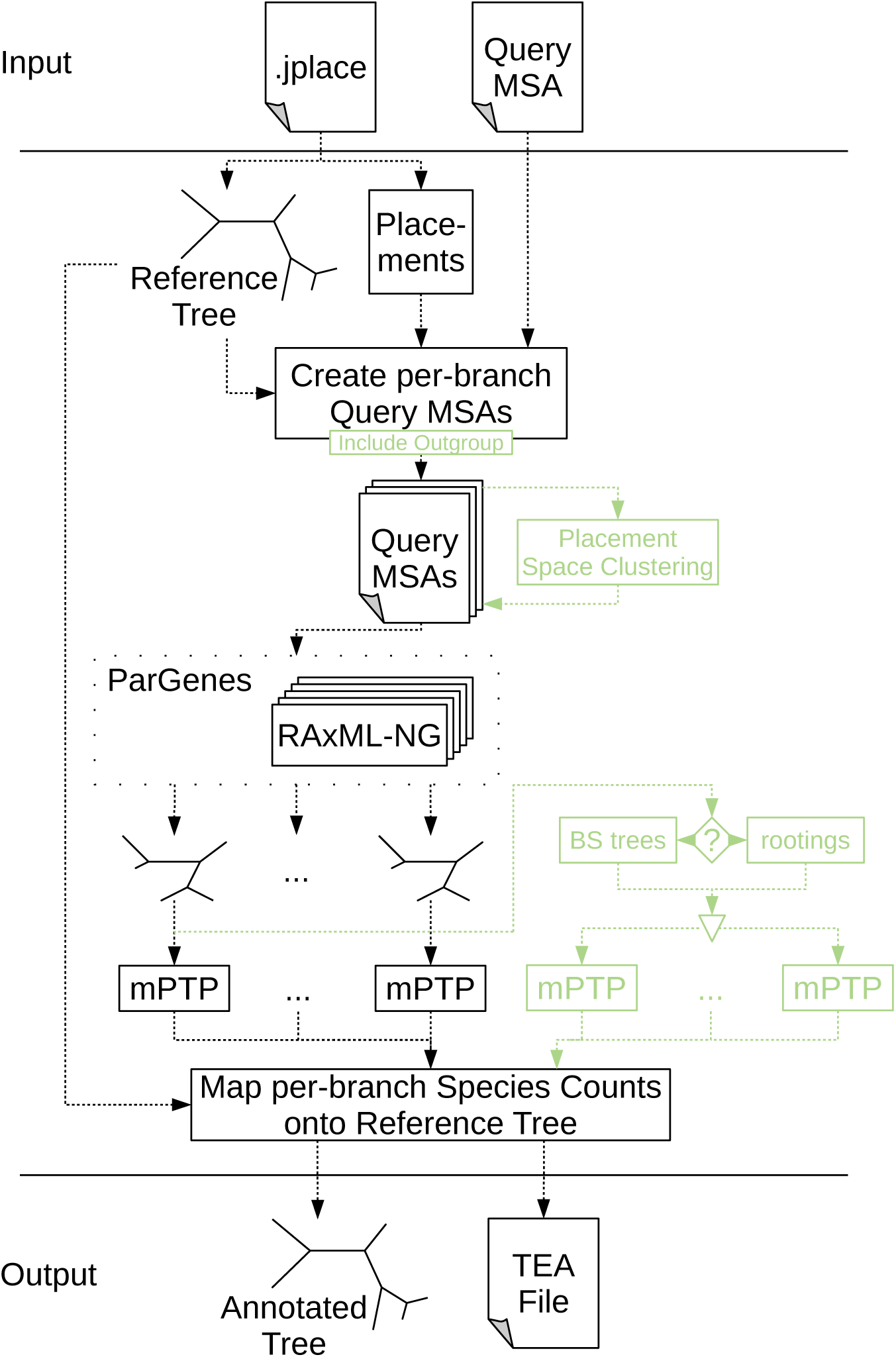
Overview of the major components of the `SCRAPP` pipeline. In green, we highlight optional components (inclusion of reference sequences for BQPs outgroup rooting, *placement space clustering* for limiting computational effort, bootstrapping or re-rooting for delimitation variance assessment).

If the number of QS in a given branch QS MSA exceeds a user-specified maximum (500 by default), we reduce the number of QS to that maximum using the two-stage clustering method described in Section 2.1. This option is necessary to maintain BQP tree inference times within reasonable limits. On empirical datasets, specific reference branches can contain more than 100, 000 QS, hence yielding the inference of a BQP computationally challenging.

Once the query MSAs have been generated for all branches of the reference tree, we infer a phylogeny for each of them separately using `RAxML-NG` (Kozlov et al., 2019). As there may be a large number of trees (potentially as many as there are branches in the reference tree) with highly variable sizes to infer, we use `ParGenes` (Morel et al., 2018) to orchestrate this tree inference process in a parallel, scalable, and efficient way. The inferred BQPs are then processed using `mPTP` to obtain a species delimitation, and corresponding species count. The information produced by each `mPTP` run is tracked for each branch that contains QS in the reference tree.

The set of inferred BQPs can optionally be expanded to calculate species count variance. Two options are available to calculate this variance: *rootings*, generates a tree set on each BQPs by enumerating all possible rootings for the unrooted BQP, or *bootstraps*, generates a given number (20 by default) of bootstrapped branch QS MSAs and then re-optimizes the branch lengths on the original BQP for each of the bootstrapped branch QS MSAs. When using these expanded BQP sets, we calculate the final species count as median over all per-branch species delimitation results (i.e., over all rootings or all bootstrap replicates).

The *rootings* and *bootstraps* options constitute two of the three principal operating modes of `SCRAPP`. The third operating mode, the *outgroup* mode, offers the rooting of the BQPs via inclusion of a reference outgroup (as described above).

Finally, `SCRAPP` generates two types of output files. Firstly, it outputs an annotated version of the reference tree in the extended NEWICK format, that can easily be visualized by a number of tree viewers (e.g., iTOL (Letunic & Bork, 2006) or Dendroscope (Huson et al., 2007)). This is useful for obtaining a high level overview of the diversity, as diversity is represented by just one species count value per reference tree branch.

To allow users to explore the results more thoroughly, for example, by inspecting the variance of the median species count, we also produce a comprehensive output file in a `json`-based file format that is analogous to the `jplace` format (Matsen et al., 2012). This format, called Tree Edge Annotations (TEA), contains the reference tree with enumerated branches, as specified in `jplace`, followed by annotation information. The annotation comprises a list of per-branch values. In `SCRAPP` this annotation includes the median species count, and the species count variance, among others. We provide a full specification and an example of the TEA format in the supplement, as well as online at https://github.com/pbdas/scrapp/wiki/TEA-format.

### 2.1 Placement space clustering

In general, phylogenetic diversity metrics face a fundamental scalability issue, as they rely on a phylogeny inferred on the QS. With increasing sequencing volumes, inferring such phylogenies under maximum likelihood becomes prohibitively expensive. Moreover, as metabarcoding/metagenomic samples typically comprise short sequences, the available signal for reliable tree inference on thousands or tens of thousands of taxa is mostly insufficient (Bininda-Emonds et al., 2000). This was the key motivation for the development of phylogenetic placement methods as a scalable and more reliable alternative.

Nonetheless, `SCRAPP` faces this same computational issue again at a different level as a reference branch may contain tens of thousands of QS. To alleviate this, we have implemented a two-stage clustering method called placement space clustering (PSC) in `SCRAPP`. PSC leverages the fact that the insertion location of a maximum likelihood placement of QS along the reference branch (the so-called proximal length) and distance from that reference branch (the so-called pendant length) can be interpreted as an embedding into a 2-dimensional euclidean space (hereafter called *placement space*). When using PSC, we map the set of placements on a branch into placement space and then perform a standard *k*-means clustering on the respective datapoints. Subsequently, we select a small number *x* of placements from each cluster as representatives of that cluster, such that *k* ∗ *x* equals the maximum desired number (as specified by the user) of sequences per branch QS MSA. More specifically, we select the top *x* := 10 sequences by number of informative (non-gap or non-undetermined) sites, thereby maximizing the potential phylogenetic signal for the subsequent tree inference.

## 3 EVALUATION

We assessed the accuracy of `SCRAPP` using both, simulated, and empirical data.

### 3.1 Simulated Data

We generated *true* species trees using the `msprime` (Kelleher et al. (2016), v0.7.3) coalescent simulator. We then used `Seq-Gen` (Rambaut and Grass (1997), v1.3.4) to generate MSAs on those trees. We generated the trees and MSAs such as to evaluate `SCRAPP` under a broad range and combination of simulation parameters (population size, mutation rates, number of taxa, etc.). In particular we investigated the influence on each parameter individually while keeping the remaining parameters fixed to a set for default values (see supplement for details).

From each simulated *true* tree and MSA, we first pruned a set of QS by removing all but one individual from each starting population. To account for incomplete reference data with lower taxon sampling density, we subsequently further pruned a given fraction (denoted as `prune fract`) of taxa uniformly at random from the trees. We then labeled the branches of the remaining reference tree by the number of query species (here assumed to be equal to the number of populations) whose *true* location is on that given branch.

We then used `EPA-ng` to place the query data back onto the tree. Next, we evaluated these placement results using `SCRAPP`, yielding an annotated NEWICK tree. Finally, we compare the reference tree with the inferred species count annotations (hereafter `SCRAPP`-tree) to the reference tree with the *true* species count annotations.

All scripts used for generating the simulated data can be found in the `SCRAPP` repository: https://github.com/Pbdas/scrapp/tree/master/simtest

### 3.2 Empirical Data

In addition to the tests on simulated data, we replicated part of the evaluation of (McCoy & Matsen IV, 2013). McCoy and Matsen IV evaluated different diversity metrics by the quality of their fit with clinical metadata, which are known from literature to correlate with alpha diversity. We chose to replicate and extend the evaluation of the Bacterial Vaginosis dataset (Srinivasan et al., 2012) (hereafter called **BV**), as we already had access to the data and the specific dataset has been particularly well studied (Czech & Stamatakis, 2019). Unfortunately, due to patient data protection issues, we cannot make this dataset publicly available. Please refer to (Srinivasan et al., 2012) and (Czech & Stamatakis, 2019) for an exhaustive exploration of the BV dataset, and a detailed description of the placement of the per sample data, respectively.

Firstly, to obtain the OTU-derived diversity measures used in the evaluation of (McCoy & Matsen IV, 2013), we performed OTU clustering using `SWARM` (Mahé et al. (2014, 2015), v3.0, `-d 1 -f`), and utilizing `VSEARCH` (Rognes et al. (2016), v2.6.2) for dereplication and filtering. We further analyzed the resulting OTU table using the R package `phyloseq` (McMurdie and Holmes (2013), v1.22.3, function `estimate richness`) to obtain the Shannon (Shannon, 1948), Simpson (Simpson, 1949), ACE (Chao & Lee, 1992), and Chao1 (Colwell & Coddington, 1994) indices.

Secondly, to assess the placement based methods, we computed a phylogenetic placement of the sample data. Note that, we did not use the reference tree given in the original publication (Srinivasan et al., 2012), as we found that the inclusion of multiple strains of the same bacterial species can produce a very flat likelihood distribution for potential placements of a single QS across individual branches of the tree (Czech & Stamatakis, 2019). Therefore, we used an appropriately modified version of the reference tree, as shown in Figure S1 in (Czech & Stamatakis, 2019). This modified reference tree only retains consensus sequences of all reference strains, such that only one taxon per species remains. The modified reference tree comprises 198 taxa.

Based on this placement data, we obtained the measures outlined in (McCoy & Matsen IV, 2013), on a per-sample basis, using the `guppy` command `fpd` (Matsen et al. (2010); McCoy and Matsen IV (2013), v1.1.alpha19-0-g807f6f3). Note that, we chose to omit the `guppy fpd --include-pendant` option to avoid overestimating diversity. The placement process does not resolve relationships between individual QS. Thus, the distance of each individual QS to the RT is denoted by a so-called pendant length. Consequently, if two or more QS are phylogenetically close to each other, but relatively distant to the RT, the common distance to the RT may be counted once per QS in the PD calculation. This can lead to potential overestimation.

Lastly, we applied `SCRAPP` to the placement data, running the analysis in the `bootstrap` operating mode, and limiting the maximum number of taxa per BQP to 1000. This again yields a `SCRAPP`-tree (see Section 3.1).

In the interest of comparability, we chose to re-implement the Balance Weighted Phylogenetic Diversity (BWPD) function using the `genesis` library (Czech et al., 2019), in a way such that it can be applied to `SCRAPP`-trees. The BWPD relies on a one-parameter function family interpolating between classical PD and a abundance weighted version of the PD. McCoy and Matsen IV chose to implement and evaluate the BWPD on placement results, which consist of precise locations and branch lengths of queries on the reference tree. In contrast to this, `SCRAPP`-trees comprise assignments of absolute numbers (species counts) to branches of the tree, without any more specific branch length information. To remedy this discrepancy, when calculating the BWPD on a `SCRAPP`-tree, we treat the species count of a branch as if it were a single placement, located at the middle of said branch, without a pendant length.

All data handling and analysis scripts used in the empirical data evaluation can be accessed online at https://github.com/Pbdas/diversity-compare.

### 3.3 Clustering and Showcase

Finally, we include a showcase test and analysis for two additional empirical datasets.

In one set of experiments, we utilize data from an study of eukaryotic community composition in neotropical soils (Mahé et al., 2017) to evaluate our PSC methodology (Section 2.1). This data is particularly challenging for phylogenetic placement, as the available reference data is too sparse to cover the diversity that was sampled. We will refer to this dataset as the **neotrop** dataset. For our purposes, we randomly selected small subsets of 1, 000 QS from this dataset and placed them on the reference tree described in (Mahé et al., 2017) (512 reference taxa). We then executed `SCRAPP` for distinct settings of `--cluster-above`, thereby limiting the maximum number of sequences per branch used in the subsequent BQP tree searches. As the randomly selected 1, 000 QS subsets produced a maximum of 298 QS placements per branch, a threshold value of 300 was selected as the benchmark against which all other runs are compared to, as this constitutes the “no clustering” case. For each clustering threshold setting and each operating mode we performed 5 independent runs of the same data in order to quantify variability introduced by the randomization component in the clustering algorithm.

In a second set of experiments, we used a large dataset from the UniEuk project (Berney et al., 2017) as a showcase for deploying `SCRAPP` on a standard parallel compute cluster. For this test, we used a phylogenetic placement of 585, 050 QS on a reference tree comprising 800 taxa, which resulted from an OTU clustering of roughly 300 million sequences (respective article in press). Unfortunately, the dataset has not yet been published, so we are yet unable to make it available. From this, `SCRAPP` identified 254, 103 QS as being placed with a LWR above the default 0.5 threshold (see Section 2). We limited the maximum number of sequences per branch to 800, and utilized the *bootstrap* operating mode, generating 100 bootstrap trees per BQP. This resulted in the inference of 1070 trees, the largest tree containing 979 taxa. `SCRAPP` further evaluated each of them via 100 distinct bootstrap MSAs.

## 4 RESULTS

For the simulated data, we calculate two distinct accuracy values. The first is the absolute difference between the inferred and the true species count on a branch in the reference tree. This absolute difference is then averaged over all branches of the reference tree that have non-zero values in either tree. We denote this accuracy metric as mean absolute per branch error (hereafter **MAE**).

More formally, let *S* and *T* be two trees with identical topologies and branch-associated values *s*_*i*_ and *t*_*i*_ for a given branch index *i*, respectively. *T* denotes the *true* tree, while *S* denotes the *SCRAPP-tree* (Section 3.1). Let *B* be the set of branch indices for which either *S* or *T* have non-zero values. We can now write the MAE as

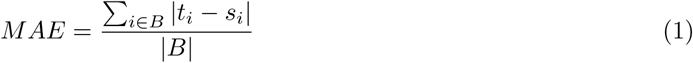

Our second accuracy metric is based on normalized per-branch species counts. For a given branch with index *k*, we calculate this normalized count based on a absolute species count *x*_*k*_ as

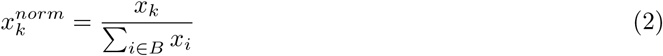

where *k* denotes the index of a given branch, and *B* is as defined above.

Further, instead of calculating the absolute difference, we calculate the relative difference:

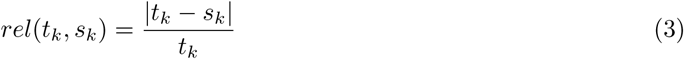

Again, *s*_*k*_ and *t*_*k*_ are the values for a given branch with index *k*, of two given trees *S* and *T* as defined above.

Finally, we again calculate the average over all relative normalized species count differences across all branches that have non-zero value, resulting in the normalized mean relative per branch error (**NMRE**).

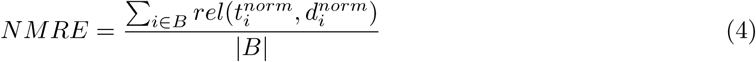

The MAE captures the deviation of the `SCRAPP`-based species count from the true species count. The NMRE quantifies the difference between the true and the inferred diversity distribution over the tree.

Regarding the accuracy evaluation of the empirical dataset, please refer to (McCoy & Matsen IV, 2013) for an in-depth description of the methods and metrics used.

### 4.1 Simulated Data

We performed a total of 270 independent simulation runs, covering all simulation space dimensions, all of their combinations with the `SCRAPP` operating modes, and repeating runs for each individual configuration 5 times. We show high level results across all runs, and stratified by operating mode, in Table 1. We observe a mean NMRE of 0.344 over all experiments. When stratified by the different operating modes, we observe the lowest overall NMRE for the rootings mode (0.3 mean NMRE).

**Table 1:** We report the mean NMRE and mean MAE, across all runs (last row) and across all runs of the specific operating modes (middle rows). *σ*^2^ denotes the variance of the given means, and CV denotes the coefficient of variation. As a reference, the mean variance among simulation replicates (identical parameter configurations but different random number seeds) was 3 × 10^*−*4^ and 3 × 10^*−*2^ for the NMRE and the MAE, respectively.

| | NMRE | $\sigma^2$ | CV | MAE | $\sigma^2$ | CV |
| --- | --- | --- | --- | --- | --- | --- |
| bootstrap | 0.356 | 0.01 | 0.27 | 5.71 | 3.00 | 0.30 |
| outgroup | 0.377 | 0.01 | 0.28 | 7.67 | 5.44 | 0.30 |
| rootings | 0.300 | 0.01 | 0.38 | 8.17 | 5.78 | 0.29 |
| across-all | 0.344 | 0.01 | 0.32 | 7.18 | 5.84 | 0.34 |

To summarize our exploration of the impact of different simulation parameters, we find that result accuracy in terms of mean NMRE increases with increasing overall population size, sample size (number of individuals per population), and sequence length, as well as decreasing `prune fract` (Section 3.1). While less pronounced, there is a trend for the NMRE to improve with increasing total tree size which may be attributed to improved taxon sampling density. This can be observed in Figure 2, which shows data for those simulation runs where we only varied the total number of starting populations (here called species). For detailed figures exploring the effect of varying individual simulation parameters, please refer to the supplement.

**Figure 2:**
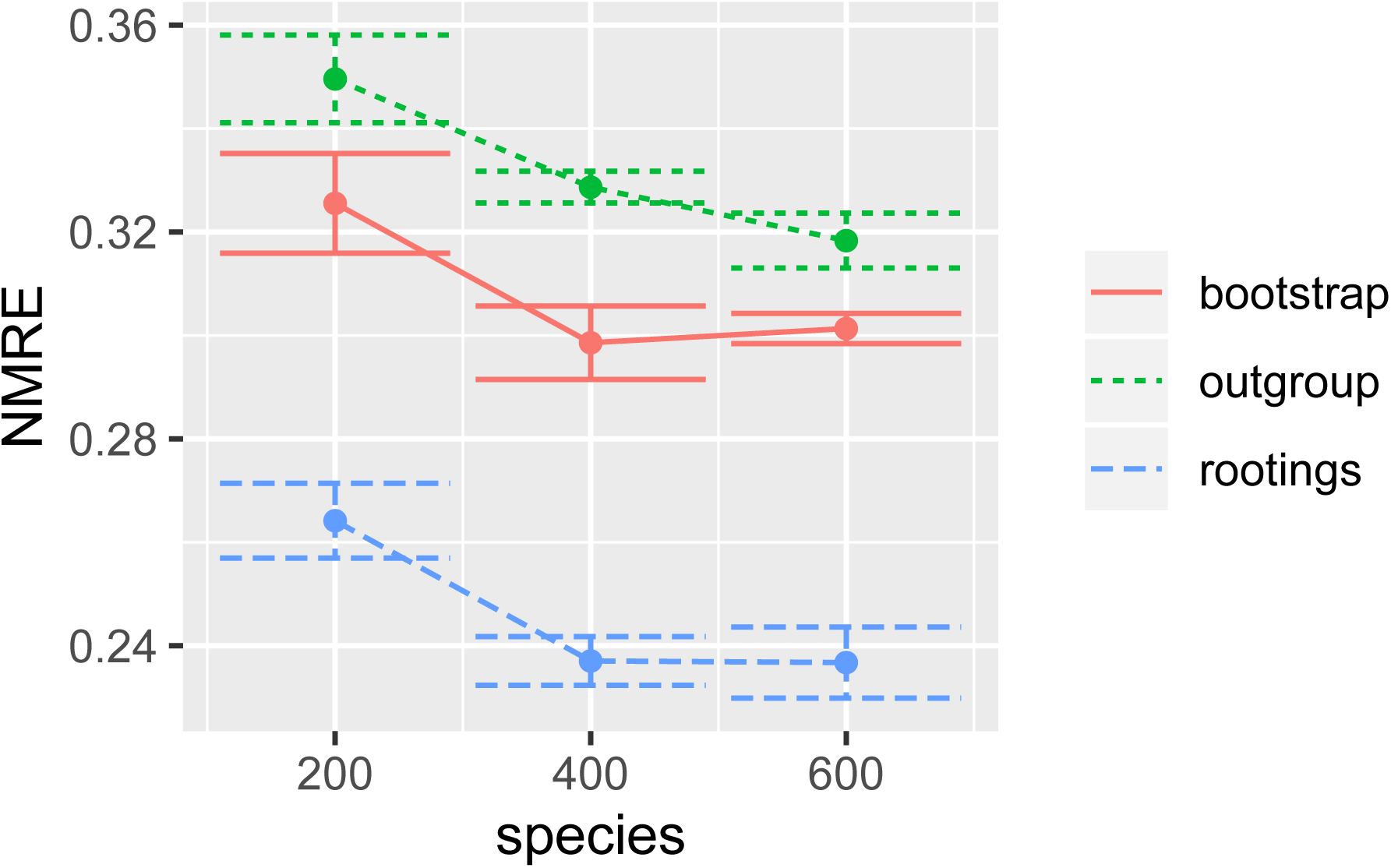
NMRE (Equation 4) for several runs on simulated datasets where we only varied the total species count of the “true” tree (the number of individual populations). Error bars denote the first standard deviation from the mean. Data was stratified by the three different operating modes of `SCRAPP` (see Section 2).

### 4.2 Empirical Data

The most important results of our evaluation based on the BV dataset are shown in Table 4.2. We were able to closely replicate the results of (McCoy & Matsen IV, 2013) (their Table 2), although we observe generally higher values for the Amsel accuracy and Nugent *R*^2^. The exception to this are the *R*^2^ values obtained from the ACE and Chao1 measures, that substantially underperform compared to the results of (McCoy & Matsen IV, 2013). As ACE and Chao1 are the only tested OTU-based metrics that specifically assign a higher weight to rare observations, we speculate that our data handling approach has reduced the number of rare OTUs. However, our results confirm the general trend that phylogenetic methods outperform OTU methods with respect to the aforementioned metrics.

**Table 2:** Correlation and predictive power of `SCRAPP` in comparison with analogous approaches on the Bacterial Vaginosis data. Amsel accuracy, Nugent *R*^2^, Amsel *p*-value, and mean rank are calculated exactly as in (McCoy & Matsen IV, 2013). Rows are sorted by mean rank. Measures suffixed by `.guppy` are calculated using guppy fpd (Matsen et al., 2010), whereas measures suffixed by `.scrapp` were calculated based on results produced by `SCRAPP`. Shannon, ACE, Chao1, and Simpson values were calculated based on an OTU clustering of the same data (see Section 3.2).

| Measure | Amsel accuracy | Nugent $R^2$ | Amsel $p$-value | Mean rank |
| --- | --- | --- | --- | --- |
| bwpd_0.25.guppy | 0.877 | 0.777 | 2.01e-35 | 2.33 |
| bwpd_0.25.scrapp | 0.874 | 0.785 | 4.02e-34 | 2.33 |
| phylo_entropy.scrapp | 0.873 | 0.782 | 4.70e-34 | 4.00 |
| bwpd_0.5.guppy | 0.873 | 0.757 | 1.03e-34 | 4.33 |
| bwpd_0.5.scrapp | 0.872 | 0.786 | 1.37e-33 | 4.67 |
| bwpd_0.scrapp | 0.873 | 0.767 | 1.49e-33 | 5.67 |
| quadratic.scrapp | 0.869 | 0.779 | 1.60e-32 | 8.33 |
| bwpd_0.75.guppy | 0.870 | 0.725 | 2.46e-33 | 9.00 |
| bwpd_0.75.scrapp | 0.868 | 0.772 | 5.10e-32 | 10.33 |
| quadratic.guppy | 0.869 | 0.718 | 7.97e-33 | 10.33 |
| bwpd_0.guppy | 0.872 | 0.713 | 2.00e-31 | 11.17 |
| unrooted_pd.guppy | 0.872 | 0.713 | 2.00e-31 | 11.17 |
| phylo_entropy.guppy | 0.869 | 0.716 | 1.43e-32 | 11.33 |
| rooted_pd.guppy | 0.871 | 0.701 | 5.73e-31 | 13.00 |
| bwpd_1.scrapp | 0.861 | 0.741 | 1.30e-29 | 13.67 |
| bwpd_1.guppy | 0.867 | 0.691 | 8.36e-32 | 14.33 |
| Shannon | 0.826 | 0.387 | 5.03e-18 | 17.00 |
| ACE | 0.822 | 0.242 | 1.41e-10 | 18.00 |
| Chao1 | 0.810 | 0.213 | 6.35e-09 | 19.00 |
| Simpson | 0.788 | 0.168 | 3.61e-08 | 20.00 |

Further, we observe a high level of agreement between metrics directly calculated from placement results, and metrics derived from `SCRAPP` results.

### 4.3 Clustering and Showcase

The results of evaluating PSC with varying clustering thresholds are shown in Figure 3. Both, the *bootstrap*, and *rootings* operating modes produced stable results, that are qualitatively similar to the tests on simulated data. However, the *outgroup* operating mode proved to be highly sensitive to the PSC, yielding high species count deviations starting at a clustering threshold of 200 (a data reduction of approx. 33%). Due to the known issues with the eukaryotic soil reference dataset at hand we hypothesize that the cause for this behavior is the sparse taxon sampling in the reference MSA. This incomplete taxon sampling induces a high branch length distance between the ingroup QS and the outgroup, as `SCRAPP` selects the phylogenetically most distant taxon in the reference tree as outgroup.

**Figure 3:**
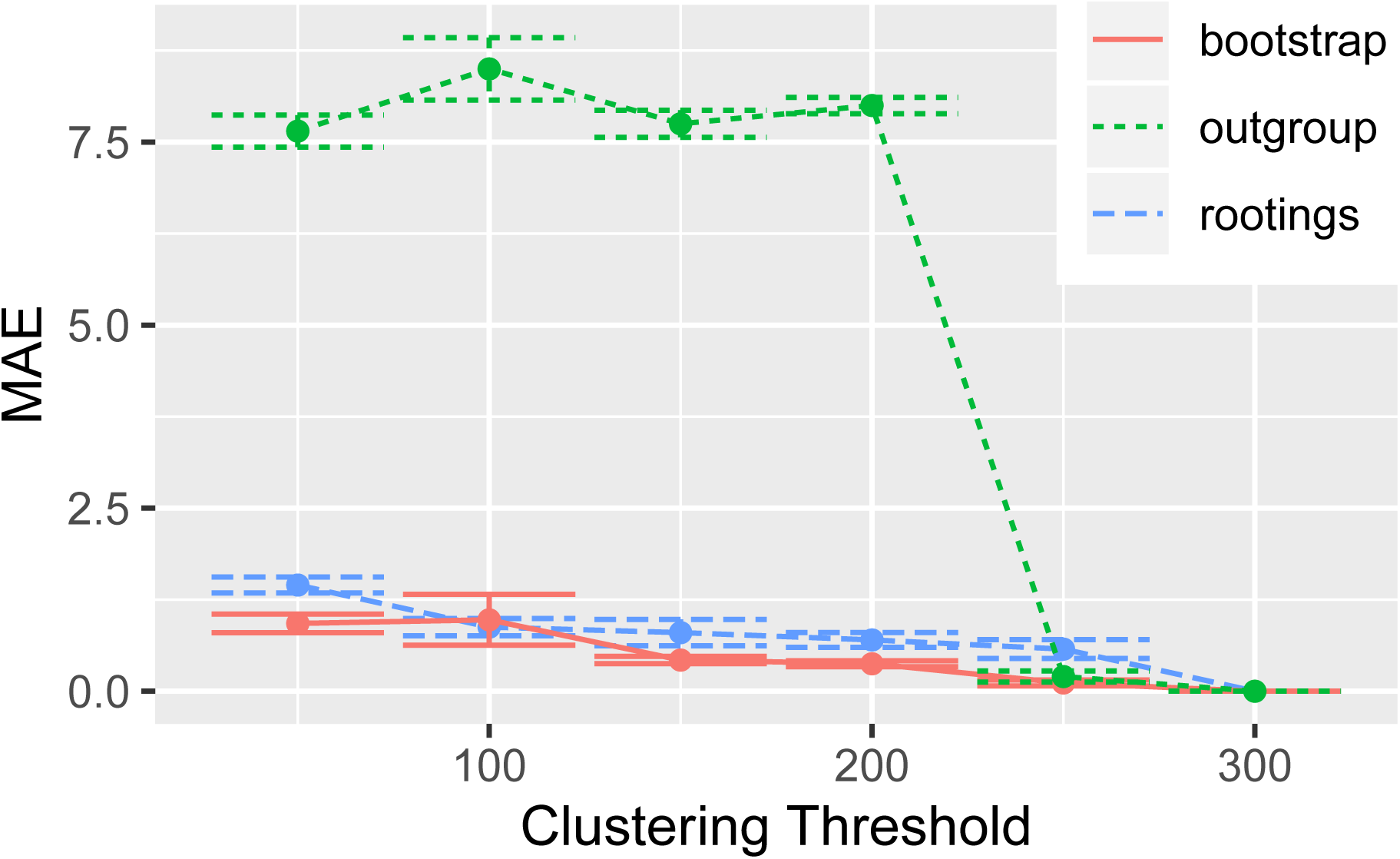
MAE (Equation 1) of multiple runs of `SCRAPP`, using different thresholds down to which placement space clustering (PSC) reduces the maximum per-branch data volume. The MAE is calculated with reference to the case of the threshold being 300, as 300 was the maximum number of QS that were placed per-branch. The underlying query and reference data are from the neotrop dataset (Section 3.3, (Mahé et al., 2017)).

As a final showcase for the scalability of `SCRAPP` on distributed computing clusters, we performed an analysis of a large dataset of 585, 050 QS placed on a 800 taxon reference tree, utilizing 50 compute nodes comprising a total of 800 cores. Running this analysis involved handling about 1 million files, of which approximately 8, 500 had to be retained as intermediate results for further downstream analysis. The total runtime under this setting was 26.5 hours, which we regard as being fast considering the size of the overall task.

## 5 CONCLUSION

We presented `SCRAPP`, a highly scalable and fully automated pipeline for diversity quantification of phylogenetic placement data. The primary goal of `SCRAPP` is to quantify the diversity distribution of a given microbial sample over the reference tree. We show that, on simulated datasets, `SCRAPP` yields phylogenetic diversity distributions with a comparatively low per-branch error rate. On empirical data, we show that alpha diversity metrics calculated on the results obtained from `SCRAPP` rank among the top of those tested in terms of predictive power, and correlation with clinical metadata.

By using MPI (Message Passing Interface), `SCRAPP` achieves a high level of parallelism, enabling the user to utilize an arbitrary number of cores in a cluster computing environment. In a selected showcase, we were able to run `SCRAPP` on a dataset with 585, 050 QS on 50 cluster nodes, using a total of 800 cores, in 26.5 hours. This run involved hundreds of tree inferences with up to 797 taxa, and the handling of approximately 1 million intermediate files.

Using placement space clustering, a novel clustering method for placements, `SCRAPP` is able to to efficiently perform dimensionality reduction of the branch QS MSA input data. This enables `SCRAPP` to tackle the scalability challenge induced by the metagenomic and metabarcoding data flood. Finally, it should be noted that issues with the underlying reference data regarding taxon sampling density may negatively affect the results when clustering is used.

## Acknowledgements

We thank S. Srinivasan and E. Matsen for providing the Bacterial Vaginosis dataset (Srinivasan et al., 2012). We extend our thanks to Paschalia Kapli and Aggelos Koropoulis for discussions and advice regarding the simulations. We thank Cédric Berney and Laura Rubinat-Ripoll for the insight that led to the development of placement space clustering. This work was financially supported by the Klaus Tschira Foundation.

## Data Accessibility

The *neotrop* dataset can be accessed via the supplementary repository to our previous publication, at https://datadryad.org/stash/dataset/doi:10.5061/dryad.kb505nc. Available in the same repository is also a anonymized version of the *BV* dataset, however no disambiguation into individual samples can be obtained from it, as this was one of the goals of patient data protection. For the same reason we are unable to share the dataset, as used in this work, publicly. The *UniEuk*-associated dataset used in the showcase is not yet published and we are unable to share it. Simulated data may be recreated from the provided scripts and instructions.

## Author Contributions

PB, LC and AS designed the software. PB and LC implemented the software. SL performed a feasibility pre-study. PB and AS wrote the paper.

## References

Barbera, P., Kozlov, A. M., Czech, L., Morel, B., Darriba, D., Flouri, T., & Stamatakis, A. (2018, 09). EPA-ng: Massively Parallel Evolutionary Placement of Genetic Sequences. Systematic Biology, 68 (2), 365–369. Retrieved from https://doi.org/10.1093/sysbio/syy054 doi: 10.1093/sysbio/syy054

Berger, S. A., Krompass, D., & Stamatakis, A. (2011). Performance, accuracy, and web server for evolutionary placement of short sequence reads under maximum likelihood. Systematic Biology, 60 (3), 291–302. doi: 10.1093/sysbio/syr010

Berney, C., Ciuprina, A., Bender, S., Brodie, J., Edgcomb, V., Kim, E., … de Vargas, C. (2017). UniEuk: Time to Speak a Common Language in Protistology! Journal of Eukaryotic Microbiology, 64 (3), 407–411. Retrieved from https://onlinelibrary.wiley.com/doi/abs/10.1111/jeu.12414 doi: 10.1111/jeu.12414

Bininda-Emonds, O. R., Brady, S., Kim, J., & Sanderson, M. J. (2000). Scaling of accuracy in extremely large phylogenetic trees. In Biocomputing 2001 (pp. 547–558). World Scientific.

Chao, A., & Lee, S.-M. (1992). Estimating the number of classes via sample coverage. Journal of the American statistical Association, 87 (417), 210–217.

Colwell, R. K., & Coddington, J. A. (1994). Estimating terrestrial biodiversity through extrapolation. Philosophical Transactions of the Royal Society of London. Series B: Biological Sciences, 345 (1311), 101–118.

Czech, L., Barbera, P., & Stamatakis, A. (2019). Genesis and Gappa: Processing, Analyzing and Visualizing Phylogenetic (Placement) Data. bioRxiv. Retrieved from https://www.biorxiv.org/content/early/2019/12/01/647958 doi: 10.1101/647958

Czech, L., & Stamatakis, A. (2019, 05). Scalable methods for analyzing and visualizing phylogenetic placement of metagenomic samples. PLOS ONE, 14 (5), 1–50. Retrieved from https://doi.org/10.1371/journal.pone.0217050 doi: 10.1371/journal.pone.0217050

Dalcin, L. D., Paz, R. R., Kler, P. A., & Cosimo, A. (2011). Parallel distributed computing using Python. Advances in Water Resources, 34 (9), 1124–1139. Retrieved from http://www.sciencedirect.com/science/article/pii/S0309170811000777 (New Computational Methods and Software Tools) doi: https://doi.org/10.1016/j.advwatres.2011.04.013

Dalcín, L., Paz, R., & Storti, M. (2005). MPI for Python. Journal of Parallel and Distributed Computing, 65 (9), 1108–1115. Retrieved from http://www.sciencedirect.com/science/article/pii/S0743731505000560 doi: https://doi.org/10.1016/j.jpdc.2005.03.010

Dalcín, L., Paz, R., Storti, M., & D’Elía, J. (2008). MPI for Python: Performance improvements and MPI-2 extensions. Journal of Parallel and Distributed Computing, 68 (5), 655–662. Retrieved from http://www.sciencedirect.com/science/article/pii/S0743731507001712 doi: https://doi.org/10.1016/j.jpdc.2007.09.005

Faith, D. P. (1992). Conservation evaluation and phylogenetic diversity. Biological Conservation, 61 (1), 1–10. Retrieved from http://www.sciencedirect.com/science/article/pii/0006320792912013 doi: https://doi.org/10.1016/0006-3207(92)91201-3

Fujisawa, T., & Barraclough, T. G. (2013, 06). Delimiting Species Using Single-Locus Data and the Generalized Mixed Yule Coalescent Approach: A Revised Method and Evaluation on Simulated Data Sets. Systematic Biology, 62 (5), 707–724. Retrieved from https://doi.org/10.1093/sysbio/syt033 doi: 10.1093/sysbio/syt033

Huson, D. H., Richter, D. C., Rausch, C., Dezulian, T., Franz, M., & Rupp, R. (2007). Dendroscope: An interactive viewer for large phylogenetic trees. BMC bioinformatics, 8 (1), 460.

Kapli, P., Lutteropp, S., Zhang, J., Kobert, K., Pavlidis, P., Stamatakis, A., & Flouri, T. (2017, 01). Multi-rate Poisson tree processes for single-locus species delimitation under maximum likelihood and Markov chain Monte Carlo. Bioinformatics, 33 (11), 1630–1638. Retrieved from https://doi.org/10.1093/bioinformatics/btx025 doi: 10.1093/bioinformatics/btx025

Kelleher, J., Etheridge, A. M., & McVean, G. (2016, 05). Efficient Coalescent Simulation and Genealogical Analysis for Large Sample Sizes. PLoS Comput Biol, 12 (5), 1–22. Retrieved from http://dx.doi.org/10.1371%2Fjournal.pcbi.1004842 doi: 10.1371/journal.pcbi.1004842

Kozlov, A. M., Darriba, D., Flouri, T., Morel, B., & Stamatakis, A. (2019, 05). RAxML-NG: A fast, scalable, and user-friendly tool for maximum likelihood phylogenetic inference. Bioinformatics. Retrieved from https://doi.org/10.1093/bioinformatics/btz305 doi: 10.1093/bioinformatics/btz305

Letunic, I., & Bork, P. (2006). Interactive Tree Of Life (iTOL): an online tool for phylogenetic tree display and annotation. Bioinformatics, 23 (1), 127–128.

Lozupone, C., & Knight, R. (2005). UniFrac: a New Phylogenetic Method for Comparing Microbial Communities. Applied and Environmental Microbiology, 71 (12), 8228–8235. Retrieved from https://aem.asm.org/content/71/12/8228 doi: 10.1128/AEM.71.12.8228-8235.2005

Mahé, F., de Vargas, C., Bass, D., Czech, L., Stamatakis, A., Lara, E., … Dunthorn, M. (2017). Parasites dominate hyperdiverse soil protist communities in Neotropical rainforests. Nature Ecology & Evolution, 1 (4), 0091. Retrieved from https://doi.org/10.1038/s41559-017-0091 doi: 10.1038/s41559-017-0091

Mahé, F., Rognes, T., Quince, C., de Vargas, C., & Dunthorn, M. (2014, 9). Swarm: robust and fast clustering method for amplicon-based studies. PeerJ, 2, e593. Retrieved from http://dx.doi.org/10.7717/peerj.593 doi: 10.7717/peerj.593

Mahé, F., Rognes, T., Quince, C., de Vargas, C., & Dunthorn, M. (2015, 12). Swarm v2: highly-scalable and high-resolution amplicon clustering. PeerJ, 3, e1420. Retrieved from https://doi.org/10.7717/peerj.1420 doi: 10.7717/peerj.1420

Matsen, F. A., Hoffman, N. G., Gallagher, A., & Stamatakis, A. (2012, 02). A Format for Phylogenetic Placements. PLOS ONE, 7 (2), 1–4. Retrieved from https://doi.org/10.1371/journal.pone.0031009 doi: 10.1371/journal.pone.0031009

Matsen, F. A., Kodner, R. B., & Armbrust, V. E. (2010). pplacer: linear time maximum-likelihood and Bayesian phylogenetic placement of sequences onto a fixed reference tree. BioMed Central Bioinformatics, 11 (1), 1–16.

McCoy, C. O., & Matsen IV, F. A. (2013). Abundance-weighted phylogenetic diversity measures distinguish microbial community states and are robust to sampling depth. PeerJ, 1, e157.

McMurdie, P. J., & Holmes, S. (2013, 04). phyloseq: An R Package for Reproducible Interactive Analysis and Graphics of Microbiome Census Data. PLOS ONE, 8 (4), 1–11. Retrieved from https://doi.org/10.1371/journal.pone.0061217 doi: 10.1371/journal.pone.0061217

Morel, B., Kozlov, A. M., & Stamatakis, A. (2018). ParGenes: a tool for massively parallel model selection and phylogenetic tree inference on thousands of genes. bioRxiv. Retrieved from https://www.biorxiv.org/content/early/2018/07/23/373449 doi: 10.1101/373449

Rambaut, A., & Grass, N. C. (1997, 06). Seq-Gen: an application for the Monte Carlo simulation of DNA sequence evolution along phylogenetic trees. Bioinformatics, 13 (3), 235–238. Retrieved from https://doi.org/10.1093/bioinformatics/13.3.235 doi: 10.1093/bioinformatics/13.3.235

Rognes, T., Flouri, T., Nichols, B., Quince, C., & Mahé, F. (2016, October). VSEARCH: a versatile open source tool for metagenomics. PeerJ, 4, e2584. Retrieved from https://doi.org/10.7717/peerj.2584 doi: 10.7717/peerj.2584

Shannon, C. E. (1948). A Mathematical Theory of Communication. Bell System Technical Journal, 27 (3), 379–423. Retrieved from https://onlinelibrary.wiley.com/doi/abs/10.1002/j.1538-7305.1948.tb01338.x doi: 10.1002/j.1538-7305.1948.tb01338.x

Simpson, E. H. (1949). Measurement of diversity. Nature, 163 (4148), 688.

Srinivasan, S., Hoffman, N. G., Morgan, M. T., Matsen, F. A., Fiedler, T. L., Hall, R. W., … Fredricks, D. N. (2012, 06). Bacterial Communities in Women with Bacterial Vaginosis: High Resolution Phylogenetic Analyses Reveal Relationships of Microbiota to Clinical Criteria. PLoS ONE, 7 (6), 1–15. doi: 10.1371/journal.pone.0037818

Tucker, C. M., Cadotte, M. W., Carvalho, S. B., Davies, T. J., Ferrier, S., Fritz, S. A., … Mazel, F. (2017). A guide to phylogenetic metrics for conservation, community ecology and macroecology. Biological Reviews, 92 (2), 698–715. Retrieved from https://onlinelibrary.wiley.com/doi/abs/10.1111/brv.12252 doi: 10.1111/brv.12252

von Mering, C., Hugenholtz, P., Raes, J., Tringe, S. G., Doerks, T., Jensen, L. J., … Bork, P. (2007). Quantitative Phylogenetic Assessment of Microbial Communities in Diverse Environments. Science, 315 (5815), 1126–1130. doi: 10.1126/science.1133420

Yang, Z. (2015). The BPP program for species tree estimation and species delimitation. Current Zoology, 61 (5), 854–865.

Zhang, J., Kapli, P., Pavlidis, P., & Stamatakis, A. (2013, 08). A general species delimitation method with applications to phylogenetic placements. Bioinformatics, 29 (22), 2869–2876. Retrieved from https://doi.org/10.1093/bioinformatics/btt499 doi: 10.1093/bioinformatics/btt499

