## Supplement for "SCRAPP: A tool to assess the diversity of microbial samples from phylogenetic placements"

### SCRAPP Supplement

November 2019

### 1 The TEA format

#### 1.1 Intention

The TEA file format aims to provide the possibility to simply and succinctly assign an arbitrary number of human and/or machine readable values to edges/branches in a given phylogenetic tree.

#### 1.2 Description

TEA uses the JSON syntax.

A TEA file has two mandatory key-value pairs: the tree, and the list of per-edge annotations. The edge annotations are also nested within a list of samples. This allows the user to specify multiple, yet semantically distinct annotations for one phylogenetic tree in a single file.

##### 1.2.1 "tree"

This key is followed by a single string specifying a single tree in NEWICK format. The tree must include edge IDs in curly brackets, for each edge/branch. The edge ID numbering starts from zero and enumerates the edges in post-order starting at the root of the tree. If the tree is unrooted, the root is set at the top level trifurcation of the given NEWICK string. We define the order of the post-order traversal as leftmost-subtree first, from the perspective of the NEWICK string.

##### 1.2.2 "views"

This is a key-value set of individual views of the tree, where each key specifies the name, and its corresponding value is a `view` on the tree. A `view` is a container holding a collection of per-edge annotation values: the `"annotation":{...}` key-value pair.

As an example (see example below), a `view` may be the SCRAPP-based species count data from the phylogenetic placement post-analysis of a metagenomic sample.

Each `view` **must** contain an `"annotation":{...}` key-value pair.

Additionally, a `view` may contain the following optional fields: * `type`: a string describing the type of the given `view`. This may be useful for grouping similar views into one.

**"annotation"** This object contains the per-edge annotation information objects. An annotation information object is not required to be complete. That is, not all edges of the tree must have an annotation and the order within this collection does not matter.

Each per-edge object is indexed by the edge number (edge ID) the annotation refers to.

The fields and structure within an annotation object are completely free and can be specified by the user.

###### 1.2.3 "meta"

This is an optional (but highly recommended) field intended to contain information about the origin of the file. As such it contains an "invocation" field specifying the command line invocation and arguments that produced the file.

###### 1.2.4 "version"

This is a mandatory field specifying which version of the TEA format the file complies with.

##### 1.3 Example

This example shows (in redacted form) how a TEA file as produced by SCRAPP might look like. The tree is specified, along with two separate metagenomic views from which the underlying phylogentic placement information was generated. Each sample contains a list of per-edge annotations comprising the results of the species delimitation(s) with mptp. Here, "count" refers to the inferred species count.

```
{
  "tree": "((A:0.2{0},B:0.09{1}):0.7{2},C:0.5{3}){4};",

  "views": {
    "leaf_emerging": {
      "type": "",
      "annotation": {
        0:{
          "count_average": 32.1,
          "count_median": 30.0,
          "count_stdev": 14.9,
          "null_score_average": 748.93,
          "null_score_stdev": 2.27e-13,
        },
        1:{"count_average": 1, ...},
        4:{"count_average": 2, ...},
      }
    },
    "leaf_growth": {
      "type": "",
      "annotation": {
        ...
      }
    }
    ...
  },

  "meta": {
    "invocation": "./scrapp.py ..."
  },
  "version": "0.1.0",
}
```


### 2 Datasets

### 3 Simulation

The default simulation parameters are given in Table 1.

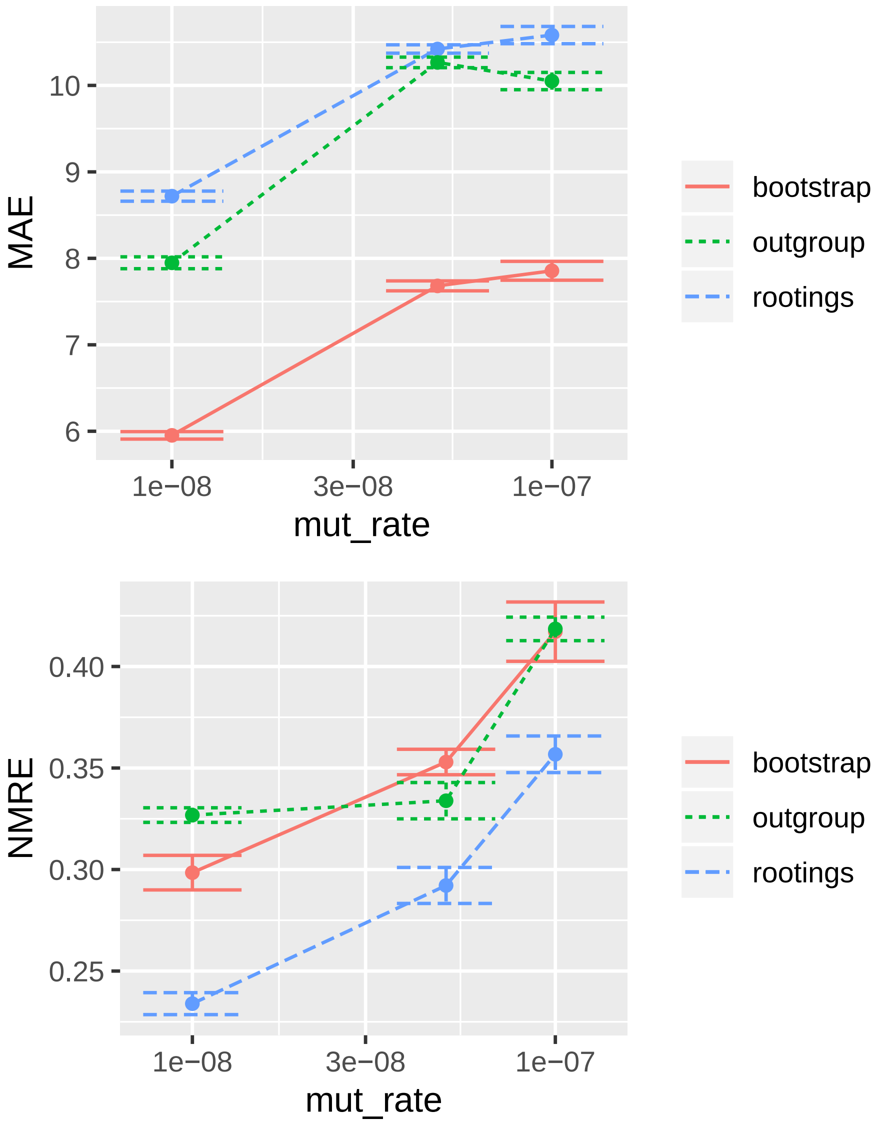

MAE
10
9
8
7
6
1e−08
3e−08
1e−07
mut_rate
bootstrap
outgroup
rootings
NMRE
0.40
0.35
0.30
0.25
1e−08
3e−08
1e−07
mut_rate
bootstrap
outgroup
rootings


Figure 1:

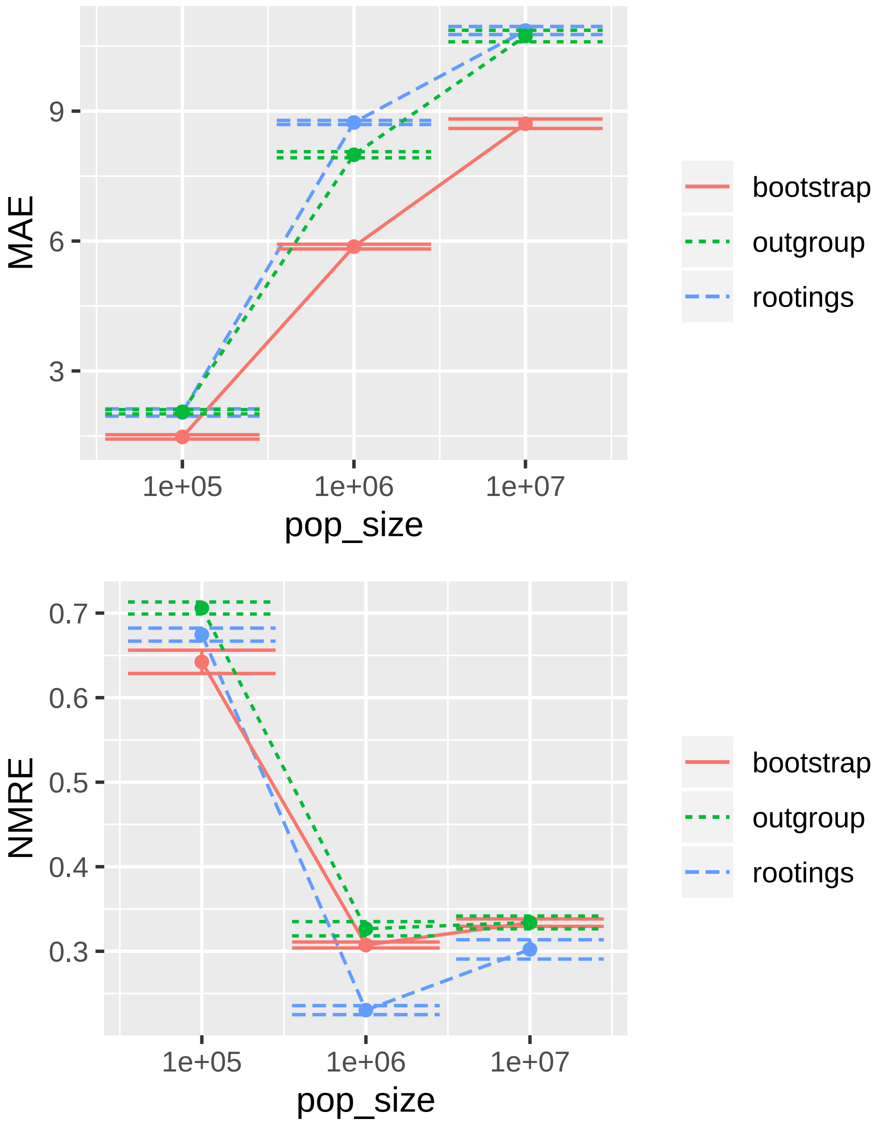

MAE
9
6
3
1e+05
1e+06
1e+07
pop_size
bootstrap
outgroup
rootings
NMRE
0.7
0.6
0.5
0.4
0.3
1e+05
1e+06
1e+07
pop_size
bootstrap
outgroup
rootings


Figure 2:

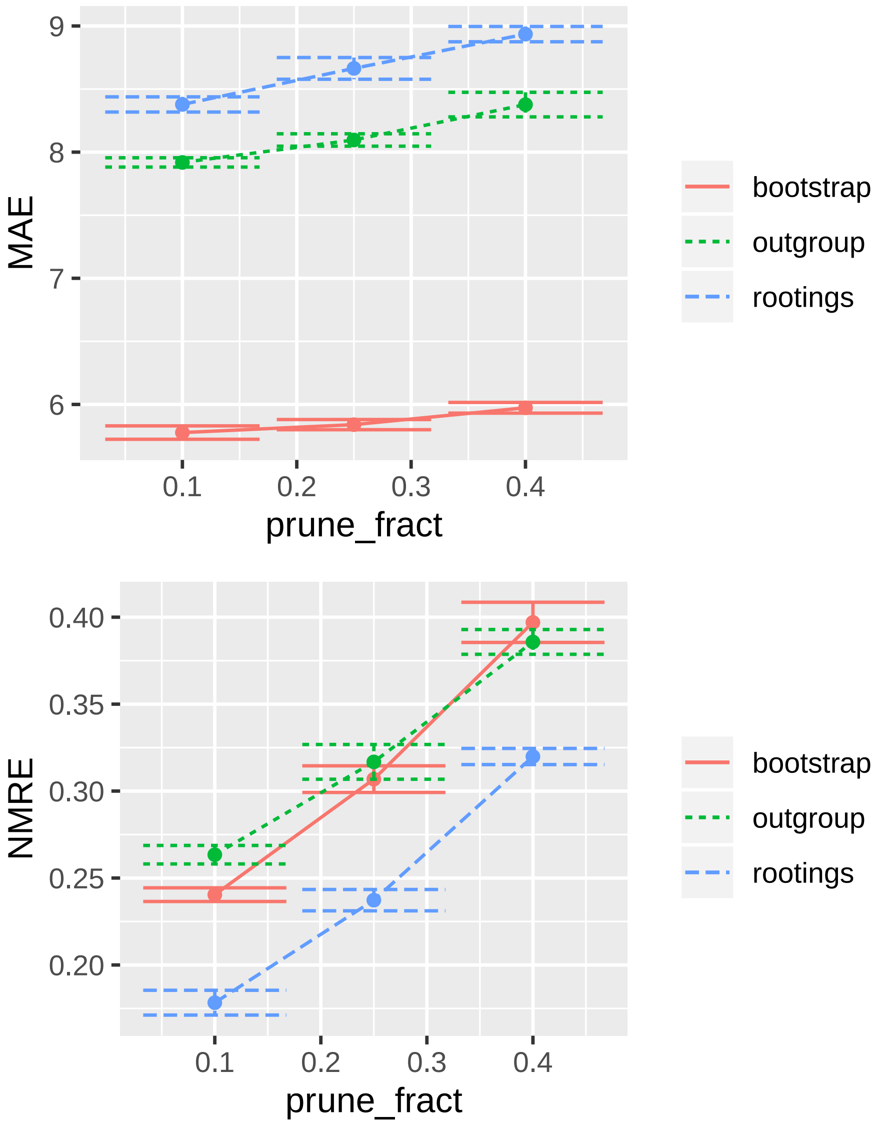
9
8
7
6
MAE
0.1
0.2
0.3
0.4
prune_fract
bootstrap
outgroup
rootings
0.40
0.35
0.30
0.25
0.20
NMRE
0.1
0.2
0.3
0.4
prune_fract
bootstrap
outgroup
rootings

Figure 3:

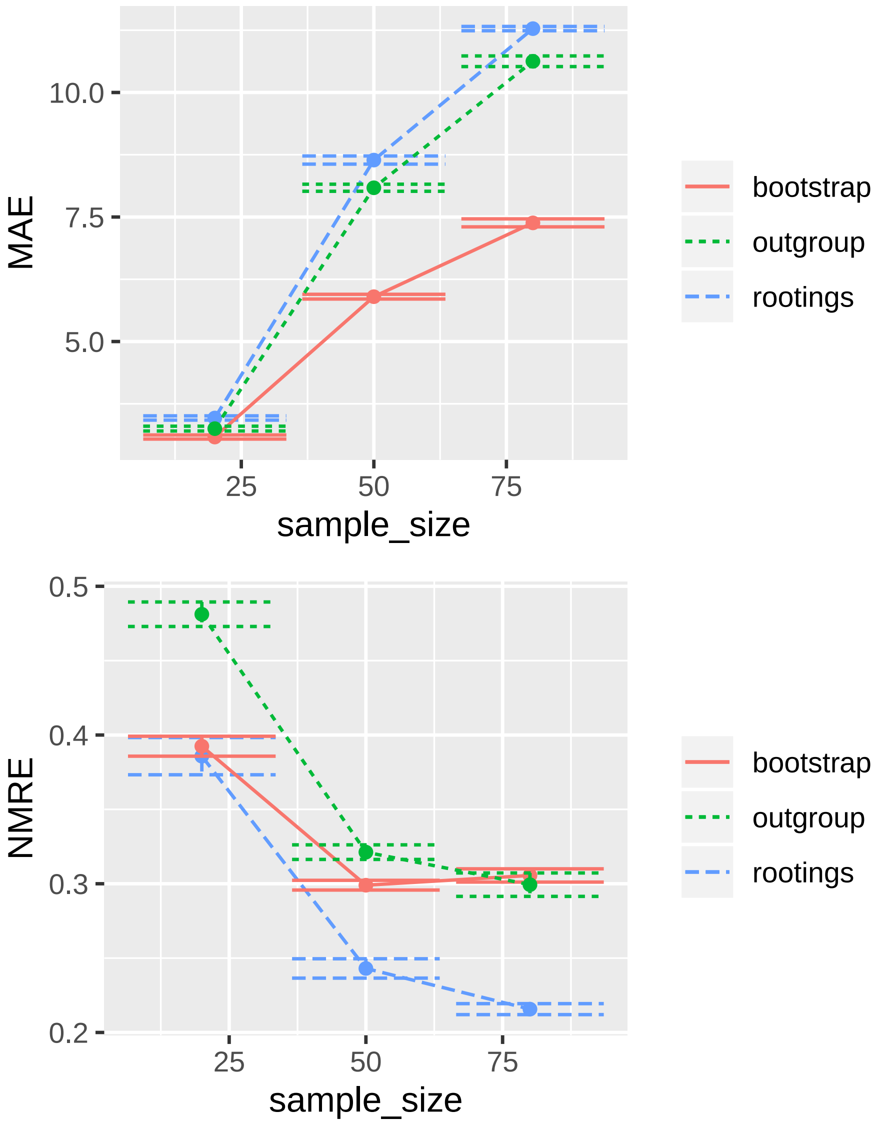

MAE
10.0
7.5
5.0
25
50
75
sample_size
bootstrap
outgroup
rootings
NMRE
0.5
0.4
0.3
0.2
25
50
75
sample_size
bootstrap
outgroup
rootings


Figure 4:

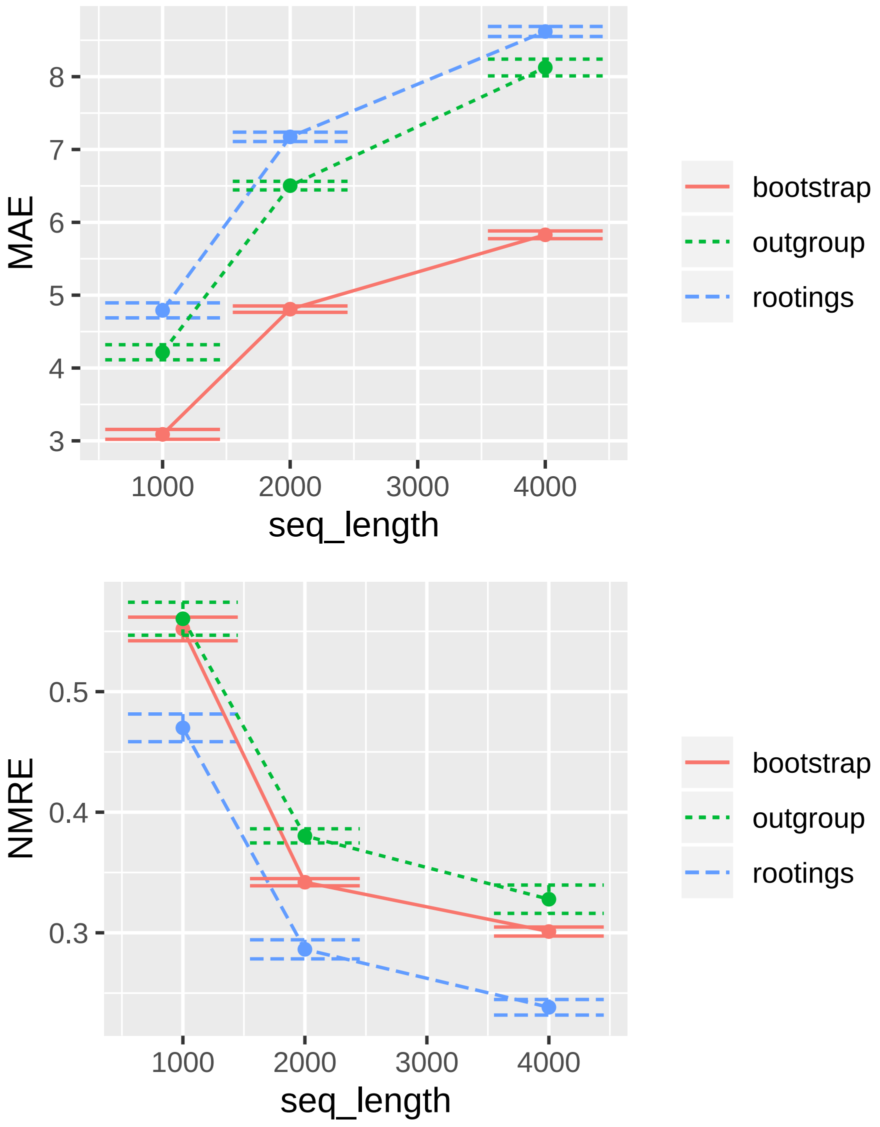

MAE
8
7
6
5
4
3
1000
2000
3000
4000
seq_length
bootstrap
outgroup
rootings
NMRE
0.5
0.4
0.3
1000
2000
3000
4000
seq_length
bootstrap
outgroup
rootings


Figure 5:

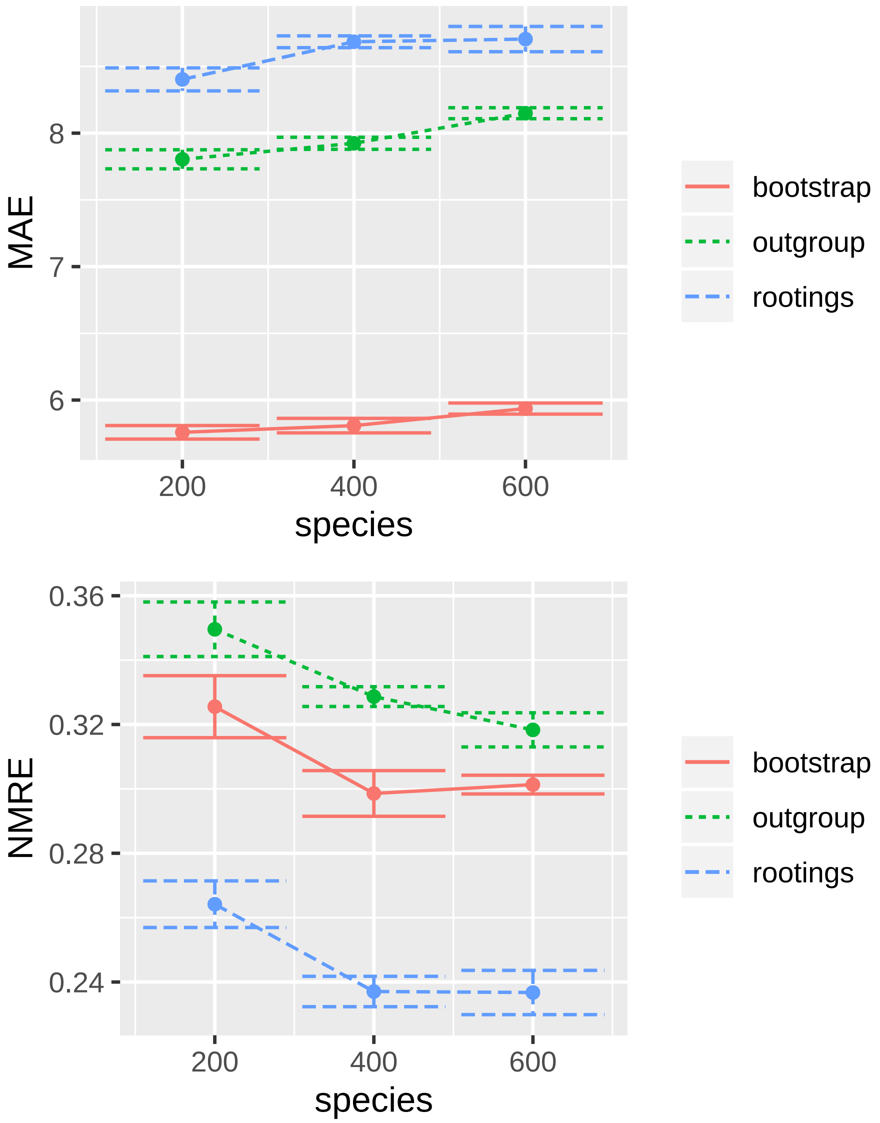
MAE
8
7
6
200
400
600
species
bootstrap
outgroup
rootings
NMRE
0.36
0.32
0.28
0.24
200
400
600
species
bootstrap
outgroup
rootings

Figure 6:

| **Parameter** | **Value** |
|---|---|
| seq_length | 4000 |
| prune_fract | 0.25 |
| pop_size | 1e6 |
| species | 400 |
| mut_rate | 1e-8 |
| sample_size | 50 |

Table 1: Default simulation parameters as used in msprime and Seq-Gen.
